# Obligate mutualistic cooperation limits evolvability

**DOI:** 10.1101/2020.11.06.371757

**Authors:** Benedikt Pauli, Leonardo Oña, Marita Hermann, Christian Kost

## Abstract

Cooperative mutualisms are widespread in nature and play fundamental roles in many ecosystems. Due to the often obligate nature of these interactions, the Darwinian fitness of the participating individuals is not only determined by the information encoded in their own genomes, but also the traits and capabilities of their corresponding interaction partners. Thus, a major outstanding question is how obligate cooperative mutualisms affect the ability of organisms to respond to environmental change with evolutionary adaptation. Here we address this issue using a mutualistic cooperation between two auxotrophic genotypes of *Escherichia coli* that reciprocally exchange costly amino acids. Amino acid-supplemented monocultures and unsupplemented cocultures were exposed to stepwise increasing concentrations of different antibiotics. This selection experiment revealed that metabolically interdependent bacteria were generally less able to adapt to environmental stress than autonomously growing strains. Moreover, obligate cooperative mutualists frequently regained metabolic autonomy, thus resulting in a collapse of the mutualistic interaction. Together, our results identify a limited evolvability as a significant evolutionary cost that individuals have to pay when entering into an obligate mutualistic cooperation.

## Introduction

Mutualistic interactions have been key for the evolution of life on earth^1^. By cooperating with members of the same or a different species, organisms gain novel capabilities without having to autonomously evolve these traits^2, 3, 4^. If the benefits resulting from these reciprocal interactions strongly outweigh the costs, significant fitness advantages can result for the interacting individuals, relative to organisms not engaging in such an interaction^5, 6^. However, an almost inevitable consequence of living within a mutualistic consortium is that both partners adapt to each other. As a result, interacting individuals frequently evolve obligate metabolic dependencies on their corresponding counterpart, eventually even losing the capacity to survive outside the interaction^7, 8, 9, 10^. By evolving such obligate metabolic dependencies, the evolutionary fate of interacting individuals is coupled. Even though it seems to be intuitively clear that living within an obligate cooperative mutualism should strongly affect the evolution of the interacting individuals, it remains generally unclear how and in which direction selective processes will be modified. Two possibilities are conceivable.

First, obligate mutualistic cooperation between two or more individuals could enhance their ability to evolve. For example, due to the usually strong mutual dependence among obligate cooperators, natural selection simultaneously acts on two different genomes. Consequently, more genetic targets are available to solve a certain evolutionary problem, thus potentially enabling mutualistic interactions to adapt faster than solitary organisms^11, 12^. Moreover, in horizontally transmitted mutualisms, the formation of new combinations among interaction partners can increase the variance among mutualistic consortia, which in turn could enhance their ability to evolve^13^. An empirical example, which is often interpreted as evidence for a positive effect of cooperative mutualisms on the evolvability of the mutualistic partners, is the rich adaptive radiation of angiosperms that resulted in the evolution of more than 300,000 species of flowering plants^14^.

Second, obligate mutualisms could also limit the potential of genotypes to respond to environmental selection pressures. Given that the ability of these cooperative interactions to survive stressful conditions likely differs between interaction partners, the individual with the lowest fitness might constrain the survival of the whole consortium (i.e. weakest link hypothesis^15^). Consequently, the niche space available to the whole association is reduced to an intersecting subset of the niche space accessible to the participating individuals^16, 17^. While some studies find indeed evidence that the survival of the whole interdependent consortium can be limited by the temperature tolerance of one of its partners^18, 19^, others report that mutualistic interactions can ameliorate environmental stress and thus even extent a species’ range limit^20^. Given that anthropogenic disturbances increasingly affect both terrestrial and aquatic habitats (e.g. climate change, pollution, etc.) and pose a threat to biodiversity on a global scale^21, 22^, a general theory of how mutualistic dependencies among individuals affects their evolutionary response to environmental stress is indispensable for predicting the broad consequences of global change for ecological communities.

However, studying how cooperative mutualisms affect the evolutionary capacity of the genotypes involved to cope with environmental stress is difficult, because the obligate nature of the interaction frequently thwarts an experimental manipulation of these system. This is why most studies so far addressed this question using comparative approaches^23, 24^ or endpoint analyses without consideration of intermediate states^25, 26^, which does not allow to infer causal relationships. The ideal way to address this question would be to subject organisms, which engage in a cooperative mutualism, to an orthogonal selection pressure and compare their ability to adapt, to the response of genetically identical individuals that live independently of their partner.

Here we take advantage of a laboratory-based model system consisting of two bacterial genotypes that previously evolved a costly cooperative mutualism (Fig. 1)^27^. Serial cocultivation of two auxotrophic strains, which were both unable to autonomously produce a certain amino acid and could only grow, when they reciprocally exchanged essential amino acids, favoured the evolution of cooperative genotypes that produced increased amounts of the traded metabolites. In contrast, amino acid-supplemented monocultures of auxotrophs, which were propagated in the same way, did not show this pattern. Positive fitness-feedbacks within multicellular clusters immediately rewarded an increased cooperative investment in either partner and could thus explain the evolution of mutualistic cooperation^27^. Importantly, the cooperative mutualism evolved in this study was highly beneficial when strains were allowed to grow in amino acid deficient environments, yet incurred significant fitness costs in the absence of these benefits.

**Figure 1.**
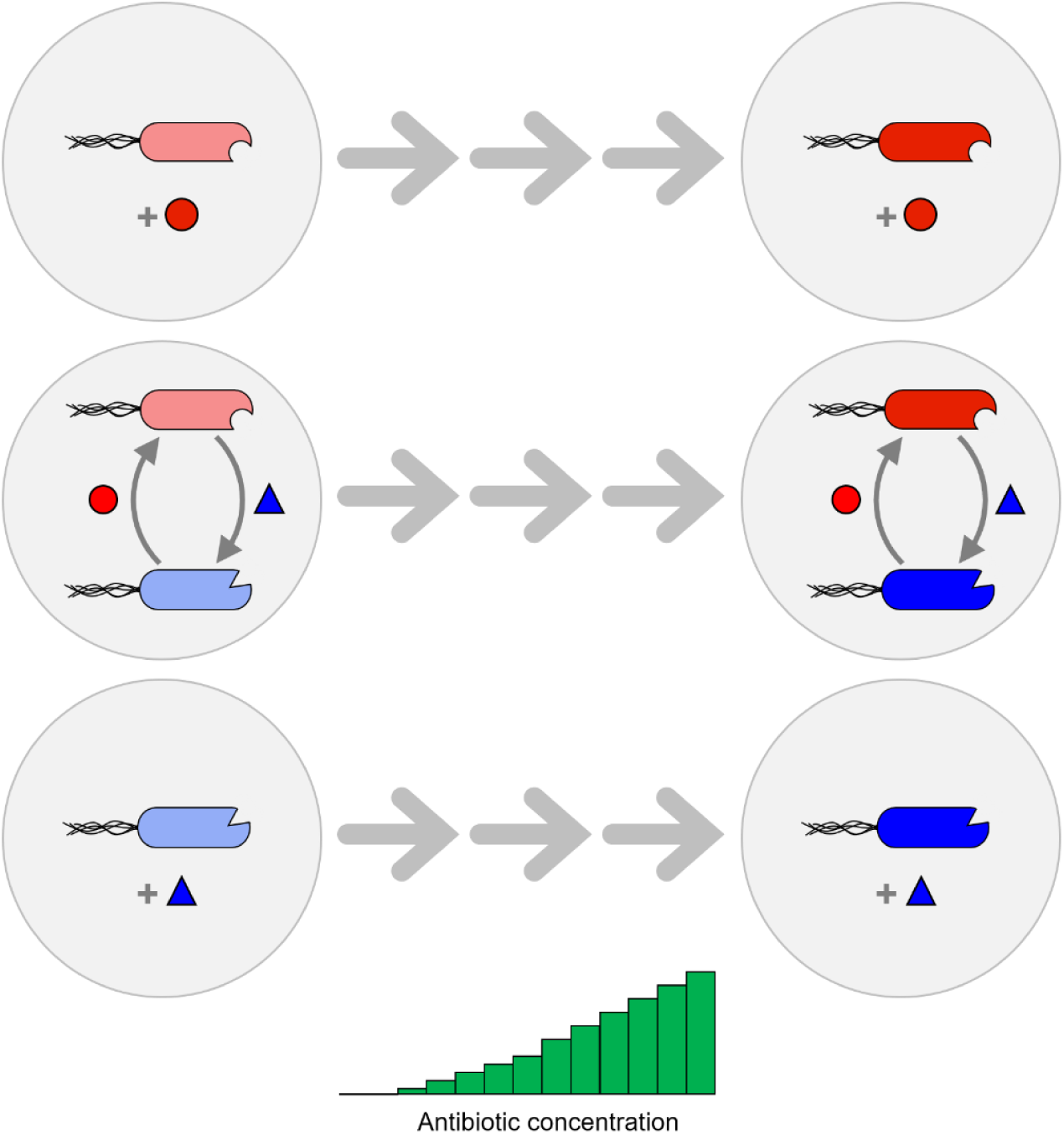
Design of the evolution experiment. In a previous study, serial coevolution of two bacterial genotypes of *Escherichia coli*, which were auxotrophic for the two amino acids tryptophan (Δ*trpB*, red cells) or tyrosine (Δ*tyrA*, blue cells), had resulted in the evolution of an obligate mutualistic cooperation between both cell types^27^. One of these mutualistic consortia was used in the current study. The two strains were either grown together in coculture (minimal medium) or in individual monocultures (minimal medium + required amino acid, 100 mM each, red circle, blue triangle). Initial populations, which were sensitive to the four different antibiotics ampicillin, kanamycin, chloramphenicol, and tetracycline, were serially propagated for 15 transfers, during which the concentration of these four antibiotics was gradually increased.

To quantitatively determine how environmental stress affects the evolvability of an obligate and cooperative mutualistic interaction, both consortia of the evolved cooperators and monocultures of the corresponding auxotrophs were subjected to gradually increasing concentrations of antibiotics (Fig. 1). The results of this experiment suggest that cooperating bacteria are more susceptible to environmental stress and are less able to adapt to these conditions compared to physiologically autonomous monocultures. Moreover, we show that albeit a synergistic coevolutionary history helped cooperative strains to deal with a current evolutionary challenge, it represented a burden when the benefit of the interaction was experimentally removed. Finally, upon exposure to high antibiotic concentrations, auxotrophic bacteria engaging in an obligate mutualism showed an increased tendency to regain metabolic autonomy and thus escape the obligate interaction. Together, our results identify a limited evolvability as a significant evolutionary cost that individuals have to pay when entering into an obligate mutualistic cooperation.

## Results

### Obligate mutualistic cooperation limits the ability of strains to adapt to environmental stress

Consortia of auxotrophic *E. coli* genotypes, which previously evolved an obligate mutualistic cooperation, were used to determine how this type of interaction affects the participating individuals’ ability to respond to environmental selection pressures. To this end, all test cultures of the current evolution experiment were serially propagated, while being subjected to a stepwise increasing concentration of one of four different antibiotics (i.e. ampicillin, kanamycin, chloramphenicol, and tetracycline) (Fig. 1). These four antibiotics differed in their mode of action. In this way, not just the effect of a single stressor was probed, but rather the more general ability of mutualistic consortia to adapt to environmental stress.

Analysing changes in population densities (OD_600nm_) of both mono- and cocultures over the course of the evolution experiment indicated that the presence of antibiotics in the growth environment had a stronger growth-reducing effect on obligate mutualistic cocultures than on monoculture controls, which were capable of independent growth (Fig. 2A,B, Supplementary Fig. 1A,B). The only exception to this pattern was the monoculture of the tyrosine auxotroph that nearly went extinct upon treatment with ampicillin (Supplementary Fig. 1C).

**Figure 2.**
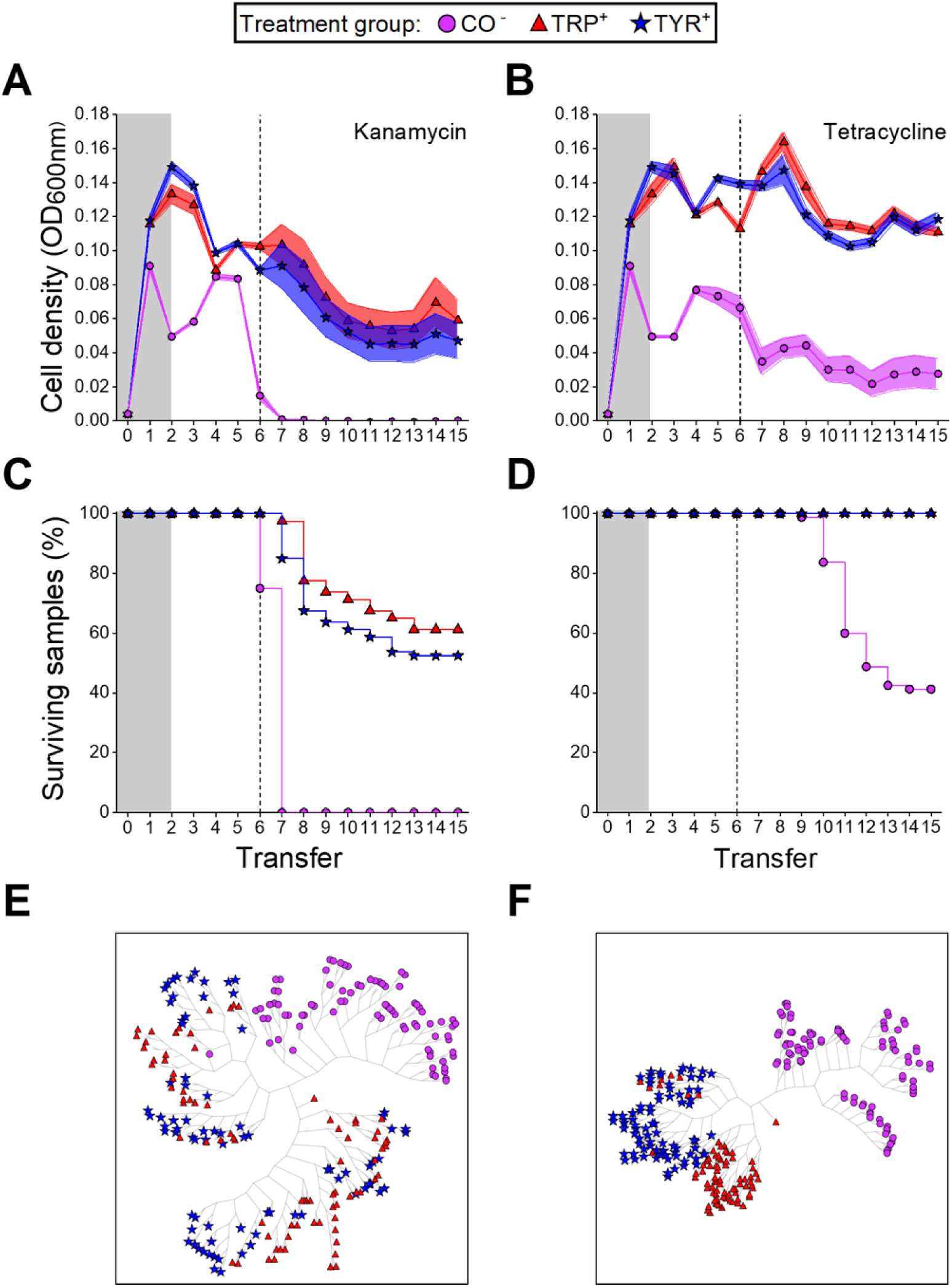
Mutualistic cooperation limits the ability of strains to adapt to environmental stress. **(A, B)** Mean growth (±95% confidence interval, n = 80 per point) quantified as OD_600nm_ and **(C, D)** proportion of surviving replicates in percent (n = 80 per strain) of auxotrophic monocultures (TRP, TYR) and mutualistic cocultures (CO) of the tryptophan (TRP) and tyrosine (TYR) auxotrophic strains over the course of the evolution experiment. Antibiotic concentrations were increased in a stepwise manner after each transfer (i.e. every 72 h) (Supplementary Fig. 3). Grey-shaded areas indicate periods without antibiotic treatment. **(A, C, E)** kanamycin treatment **(B, D, F)** tetracycline treatment. Dashed lines mark the point, at which antibiotic concentrations exceeded sub-MIC values. Monocultures were supplemented with both amino acids (100 mM each), while cocultured bacteria depended on the amino acids provided by their respective cross-feeding partner. **(E, F)** Clustering trees of cell density profiles of the different cultures across transfers in the evolution experiment. Each leaf within a given tree represents a replicate (n = 80 per strain). A radial embedding layout was used to display trees.

To further analyse differences between treatment groups, the survival of cultures in the evolution experiment was compared by applying log-rank tests for each pair of cultures. This test revealed significant differences between mutualistic consortia and monoculture controls (log-rank test: P < 0.001, Fig. 2C,D, Supplementary Fig. 1C,D, Supplementary Table 1), suggesting that mutualistic cocultures were more likely to go extinct than monocultures of auxotrophs. Upon reaching sub-MIC levels after the sixth transfer, a clear difference between the two bactericidal antibiotics (ampicillin, kanamycin) and the bacteriostatic agents (chloramphenicol, tetracycline) emerged. While bactericidal antibiotics drove all cooperative cocultures to extinction (Fig. 2C, Supplementary Fig. 1C), a subset of all cultures treated with bacteriostatic agents survived until the end of the antibiotics ramping experiment (Fig. 2D, Supplementary Fig. 1D). In all four antibiotics used, survival distributions of cocultures showed a significantly increased death rate compared to monocultures of both tryptophan- and tyrosine-auxotrophic genotypes (log-rank test: P < 0.001, Fig. 2C,D, Supplementary Fig. 1C,D, Supplementary Table 1). When monocultures were statistically compared, a significant difference was only observed for ampicillin-treated cultures, but not the groups treated with the other three antibiotics (log-rank test: P < 0.001, Supplementary Fig. 1C, Supplementary Table 1).

Finally, an unsupervised learning algorithm was applied to identify differences and similarities in the evolutionary trajectories of monocultures and cocultures. This analysis revealed in all cases the emergence of clusters that almost exclusively consisted of coculture replicates (Monte-Carlo resampling after n = 10^6^ permutations: P < 10^−6^, Fig. 2E,F, Supplementary Fig. 1E,F). This observation suggests cocultures followed a distinct evolutionary path that was significantly different from the trajectories of monoculture controls. For the monocultures treated with kanamycin, chloramphenicol, and tetracycline, the algorithm consistently detected clusters composed of a mixture of tryptophan and tyrosine auxotrophic monocultures (Fig. 4E,F, Supplementary Fig. 1F), while in the case of ampicillin, a clearly separated set of two clusters emerged (Monte-Carlo resampling after n = 10^6^ permutations: P < 10^−6^, Supplementary Fig. 1E). Taken together, these results clearly show that mutualistic cooperation limits the ability of obligate mutualisms to adapt to environmental selection pressures.

### Strain-level differences cause the increased susceptibility of cooperative consortia to environmental stress

Next, we asked whether or not the reduced ability of cooperative consortia to cope with antibiotic-mediated selection was due to differences among individual genotypes. To address this issue, the minimal inhibitory concentration (MIC) of the focal chloramphenicol- or tetracycline-treated (i.e. the two bacteriostatic antibiotics) mono- and cocultures was determined and compared among groups.

The results of this analysis revealed that the resistance level reached by coevolved consortia was significantly lower than the one of monoevolved controls (Tamhane’s post hoc test: P < 0.05, Fig. 3, Supplementary Table 2). The only exception to this was the case of the monoevolved tyrosine auxotroph, whose MIC for tetracycline did not differ significantly from the values reached by the corresponding cocultures (Fig. 3B). Strikingly, comparing resistance levels achieved by monocultures of auxotrophs that evolved as part of a mutualistic consortium to the genotypes that were cultivated as monocultures, revealed that coevolved strains reached a significantly lower MIC than the corresponding monocultures (Fig. 3). Specifically, in the case of the tetracycline-treated populations, the two auxotrophic strains showed a consistent difference between both evolutionary backgrounds, which, in combination, can explain the reduced resistance levels that the respective coevolved consortia reached when treated with this antibiotic (Fig. 3B). Together, these results demonstrate that differences in the ability of individual strains to adapt to environmental stress limited the survival and thus evolvability of the entire consortium.

**Figure 3.**
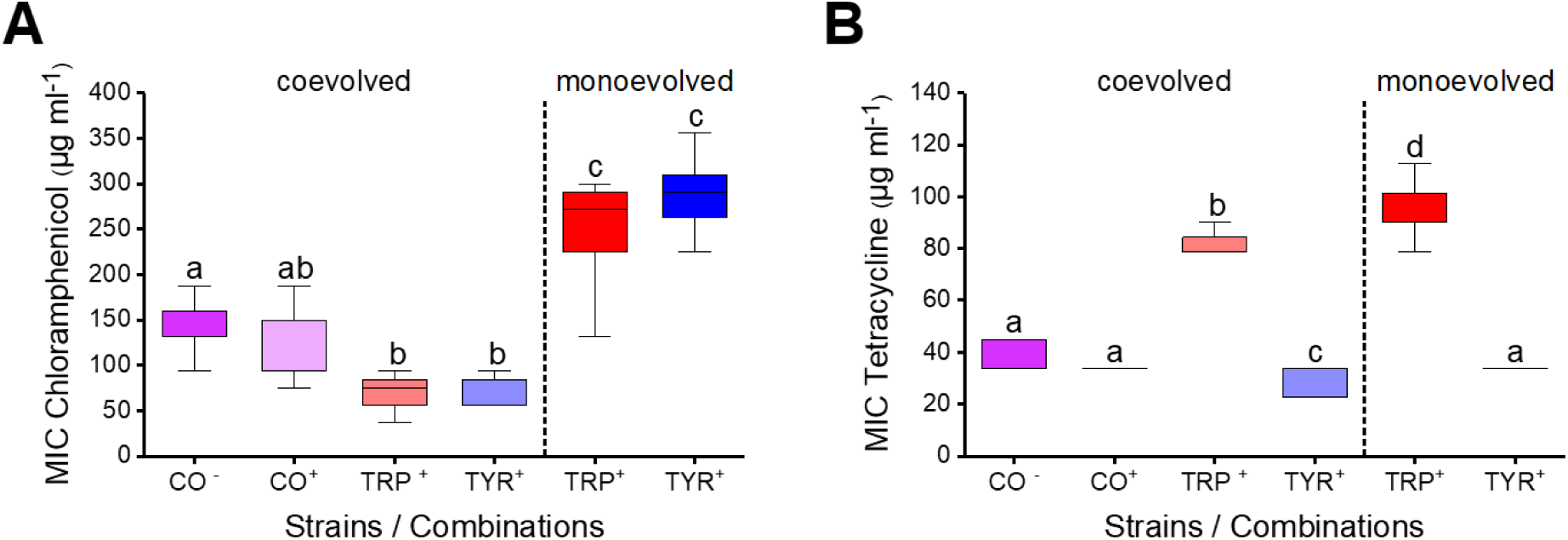
Strain-level differences cause increased susceptibility of cooperative consortia to environmental stress. The resistance levels of evolved strains treated with **(A)** chloramphenicol and **(B)** tetracycline were assessed by determining the MIC values of the corresponding derived strains. Cocultures (CO) and monocultures of the tryptophan (TRP) and tyrosine auxotrophs (TYR) from the different regimes with (^+^) and without (^-^) amino acid supplementation are displayed. Box plots show median values (horizontal line in boxes) and the upper and lower quartiles (i.e. 25-75% of data, boxes). Whiskers indicate the 1.5x interquartile range. Different letters indicate significant differences between strains (ANOVA followed by Tamhane’s post hoc test: P < 0.05, n = 8).

### Previous co-adaptation to environmental stress limits the evolvability of mutualistic consortia

Given that obligate metabolic cooperation can constrain adaptive evolution (Figs. 2 and 3), we asked how a previous coevolutionary history in a stressful environment affects the ability of a mutualistic consortium to adapt to the same environmental challenge. We hypothesized that a shared coevolutionary history should enhance the ability of derived cocultures to cope with a previously experienced environmental stress when individuals interact with each other, yet limit their ability to tolerate increased stress levels when the otherwise obligate interaction is (experimentally) uncoupled. To test this hypothesis, pairwise consortia consisting of either the two coevolved auxotrophs or the two corresponding monoevolved genotypes that have been previously exposed to the two antibiotics tetracycline and chloramphenicol were again subjected to continuously increasing concentrations of the same two antibiotics. This time, however, the experiment was performed by cultivating both coevolved and monoevolved strains as cocultures in both the absence and presence of environmentally-supplied amino acids. This experimental design allowed to experimentally disentangle the effect of a shared coevolutionary history (i.e. coevolved versus monoevolved) from effects emanating from the interaction itself (i.e. with versus without environmentally-supplied amino acids).

This experiment showed that in the absence of amino acid supplementation, coevolved cocultures of obligately cooperating genotypes were significantly better able to cope with the antibiotic to which they have been previously exposed (linear mixed model for chloramphenicol and tetracycline: P < 0.001, Fig. 4A,C, Supplementary Table 3) than cocultures of strains that previously had adapted individually to the corresponding antibiotic (CO_AUX_, Supplementary Table 4). However, when the obligate dependence between bacterial partners was relaxed by externally providing the required amino acids, the observed pattern changed to the opposite. Under these conditions, individually evolved strains reached significantly higher population densities than coevolved strains (linear mixed model for chloramphenicol and tetracycline: P < 0.001, Fig. 4B,D, Supplementary Table 3). In other words, coevolved cooperators were better off when survival depended on a metabolic interaction between both strains, while monoevolved strains had an advantage when the need to interact was experimentally removed.

**Figure 4.**
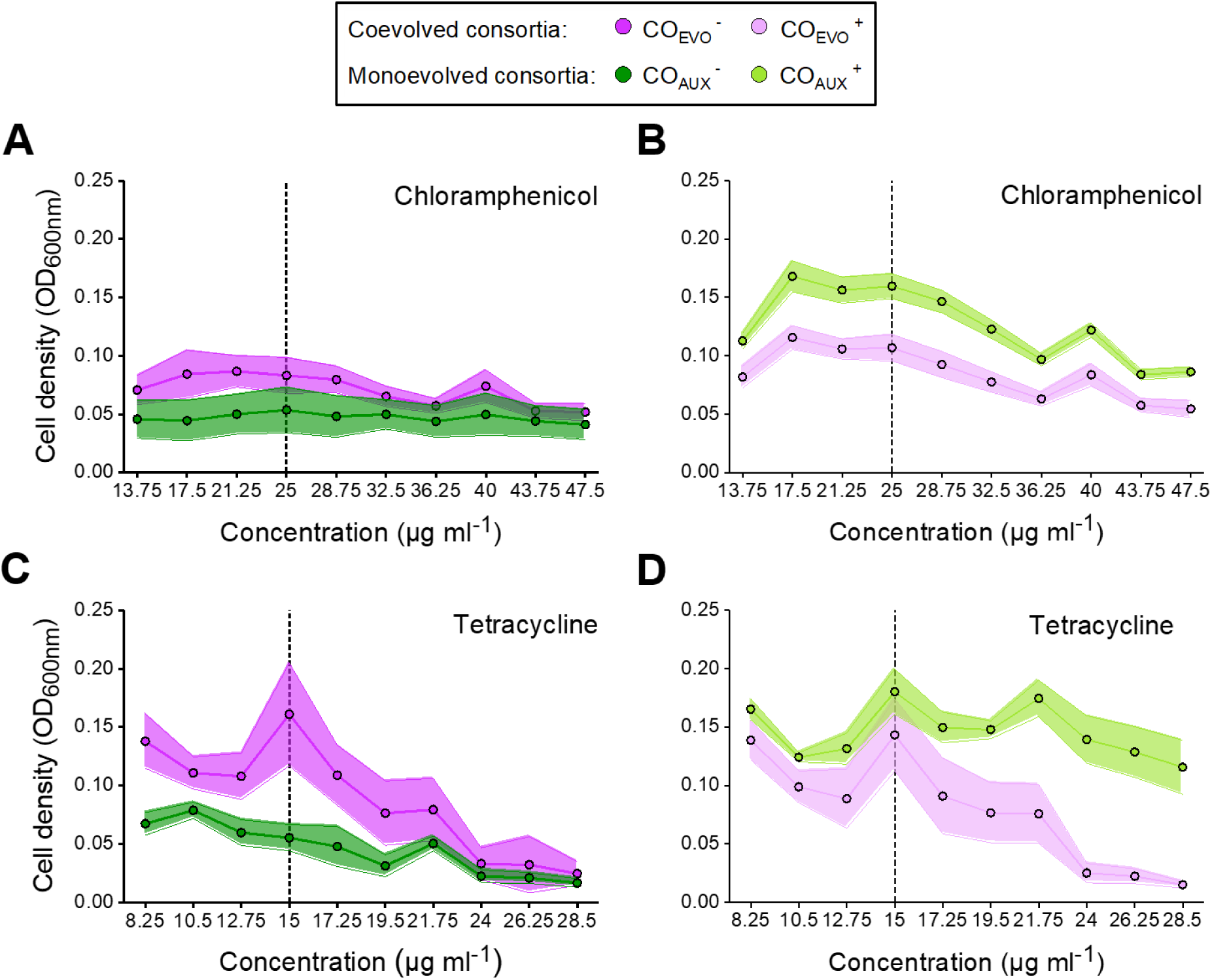
Both coevolutionary history and mutualistic dependence affect the ability of cooperative mutualists to adapt to environmental changes. Shown is the mean growth (±95% confidence interval) determined as population density (OD_600nm_) of coevolved auxotrophs (CO_EVO_) and cocultured monoevolved auxotrophs (CO_AUX_). All cultures were grown **(A, C)** with supplementation of tryptophan and tyrosine (100 mM each) or **(B, D)** without amino acid supplementation. Populations were treated with **(A, B)** chloramphenicol or **(C, D)** tetracycline. The dashed lines mark the typical working concentrations of the respective antibiotic used to select for resistant strains. To compare between strain types in the presence or absence of amino acids, contrasts between growth levels were calculated (P < 0.05, n = 8 per point).

Statistically comparing the growth response of consortia of coevolved genotypes when cultivated in the absence and presence of amino acids suggested that coevolved bacteria did not benefit from an external supplementation with amino acids (linear mixed model for chloramphenicol: P = 0.067 and for tetracycline P = 0.74, Fig. 4, Supplementary Table 3). In contrast, the growth of individually-evolved auxotrophs in the presence of elevated antibiotic concentrations strongly increased upon amino acid supplementation (linear mixed model for chloramphenicol and tetracycline: P < 0.001, Fig. 4, Supplementary Table 3). These findings imply that coevolution curtailed the ability of strains to exist outside the interaction. This result is consistent with the interpretation that an increased dependence among genotypes coupled their evolutionary fate, thus limiting their evolvability.

### Increasing environmental stress can destabilize obligate mutualistic cooperation

In situations where it is costly to obligately interact with another individual, natural selection should favour types that evolve metabolic independence. To assess whether this also happened in the course of the antibiotics ramping experiment, the population-level proportion of mutants that evolved metabolic autonomy (i.e. reverted to a prototrophic phenotype) was quantitatively determined. For this, terminal populations of both coevolved and monoevolved auxotrophs, which have been exposed to increasing concentrations of bacteriostatic antibiotics, were randomly chosen to assess the population-level fraction of reverted phenotypes. This screening revealed indeed that a subpopulation of the initial tyrosine auxotrophic genotypes regained the ability to grow without tyrosine supplementation. Interestingly, the rate of phenotypic reversion was significantly increased in strains that evolved in coculture relative to the respective monocultures (chi-square test for chloramphenicol TYR revertants: P < 0.001, χ^2^ = 42.50, df = 1 and for tetracycline TYR revertants: P < 0.001, χ^2^ = 43.00, df = 1, Fig. 5). In contrast, no revertants were detected among 572 screened colonies that were derived from 60 populations of monocultured and cocultured tryptophan auxotrophic strains. Particularly striking was the observation that no tryptophan-auxotrophic mutants could be detected in any of the tetracycline-treated cocultures (Fig. 5). This indicates that the newly evolved phenotypic revertants either outcompeted their interaction partner or simply outnumbered them (lower detection limit: 2.5 x 10^4^ cells ml^-1^). Together, these results confirm that under conditions that increase the cost of mutualistic cooperation, natural selection will favour autonomous types that abandon the obligate interaction. As a result, cooperative interactions are lost from populations, thus favouring metabolically autonomous types.

**Figure 5.**
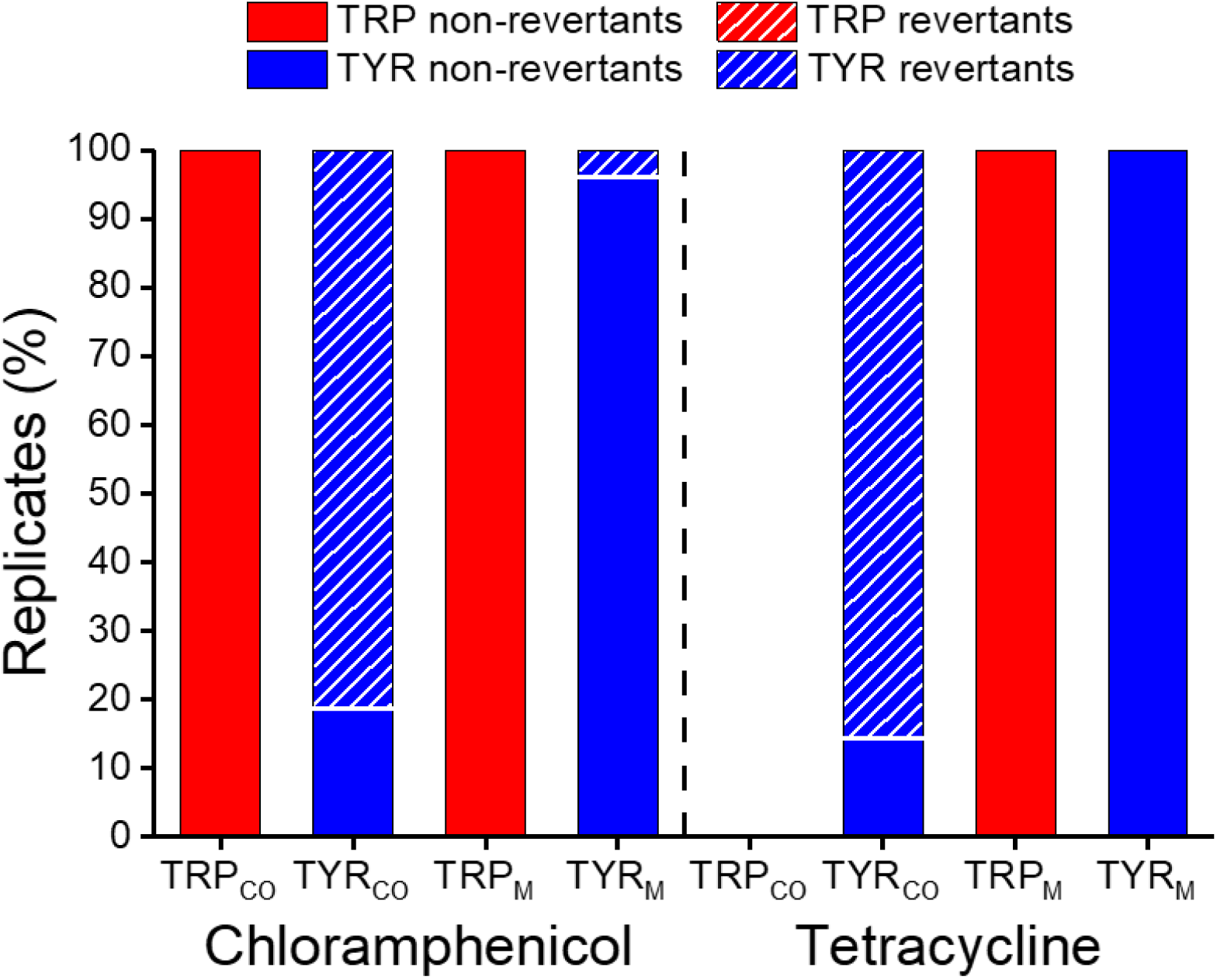
Environmental stress favours reversion to metabolic autonomy. Shown is the population-level proportion of initially auxotrophic genotypes that evolved in the presence of chloramphenicol (left) and tetracycline (right), which remained auxotrophic (filled bar) or reverted to prototrophy (hatched bars). Populations of monoevolved genotypes (TRP_M_, TYR_M_) and genotypes isolated from coevolved cultures (TRP_CO_, TYR_CO_) are compared. The plotted colony counts (%) are the number of colonies analysed per strain relative to the total number of colonies tested in the respective cultures. In both cases, reversion rates of coevolved types were significantly higher than the one of their monoevolved counterpart (chi-square test: P < 0.05. n = 243-358).

## Discussion

Mutualistic interactions significantly affect the Darwinian fitness of the organisms involved^28, 29, 30^. However, it remains largely unclear how the tight ecological coupling between both interaction partners affects their ability to adapt to environmental changes^31, 32, 33^. Here we addressed this issue using an experimental evolution approach. Specifically, two bacterial genotypes that previously evolved an obligate cooperative cross-feeding interaction were exposed to a continuously increasing concentration of one of four antibiotics. The evolutionary responses of these cocultures were compared to those of monoculture of the same two genotypes, which were cultivated by providing them with the required amino acids. The results of these experiments revealed that (i) individuals participating in an obligate mutualistic cooperation are less able to adapt to environmental stress than metabolically independent organisms (Figs. 2, 3 Supplementary Fig. 1), (ii) coevolution restrains the ability of interaction partners to respond to changing environmental selection pressures (Fig. 4), and (iii) stressful environments can select against obligate mutualisms and favour the evolution of metabolic autonomy (Fig. 5).

The main aim of this study was to distinguish between two competing hypotheses on how being part of an obligate cooperative mutualism affects the evolutionary potential of the organisms involved. First, due to the previously reported enhanced rate of evolution^27, 34^, synergistically interacting bacteria could be better able to adapt to stressful environments. In addition, members of a mutualistic consortium might directly benefit from adaptations of their corresponding partner. Alternatively, engaging in an obligate cooperative mutualism could constrain the ability of the participating individuals to respond to environmental change with evolutionary adaptation^35^. The presented experimental evidence clearly supported the second hypothesis.

In our study, cocultures of mutualistically interacting *E. coli* strains generally reached lower population densities and reduced resistance levels as compared to the corresponding monoculture controls (Figs. 2, 3, and Supplementary Fig. 1). This finding is surprising, given that several previously published studies report about so-called cross-protection mutualisms, in which resistant strains protect co-occurring sensitive strains in their environment, thus allowing them to grow in the presence of antibiotics^30, 36, 37^. In particular, the close physical contact among interacting individuals of the focal model system analysed in this study should have facilitated cross-protection. However, one main difference to these previous studies, which found evidence for cross-protection, is that the corresponding experiments were already initiated with antibiotic resistant genotypes^30^. In contrast, all strains used in our study were initially sensitive to all antibiotics and, thus, had to evolve resistance *de novo*. Additionally, our results did not support the *weakest link* hypothesis proposed by Adamowicz *et al*. (2018). The only case, in which a behaviour matching the predictions of this hypothesis was observed in our study, was the one of ampicillin-treated cultures (Supplementary Fig. 1A). In this case, both cocultures and monocultures of the tyrosine auxotroph went extinct almost simultaneously, suggesting that the pairwise consortium might have collapsed due to the lower resistance levels of the tyrosine auxotroph. However, both the survival analysis and the analysis of the clustering tree clearly showed that the ampicillin-treated coculture and the tyrosine auxotrophic strain clearly followed a different evolutionary trajectory (Supplementary Fig. 1C,E). Furthermore, it was found that the coevolved cocultures, which have been treated with bacteriostatic antibiotics, were generally more resistant to the focal antibiotic (chloramphenicol or tetracycline) than at least one of the two partners grown in monoculture (Fig. 3 ‘coevolved’). This finding is in contrast to expectations of the ‘weakest link’ hypothesis, which predicts that the antibiotic tolerance of a given consortium should be determined by the member with the lowest tolerance^15^. Together, these results clarify that the ‘weakest link’ hypothesis is likely not the main factor explaining the observed evolutionary patterns.

In our experiments, mutualistic cocultures were generally less able to adapt to environmental change than metabolically autonomous monocultures (Fig. 2, Supplementary Fig. 1). One explanation for these differences could be the tight ecological coupling that evolved within consortia of mutualistically interacting bacteria. The combination of strains used here was derived from a previous study, in which two auxotrophic mutants evolved a mutualistic cooperation at a cost to themselves^27^. The resulting strains showed a reduced ability to utilize environmental amino acids and rather relied on a supply of these essential nutrients from their corresponding partner. This rather extreme form of physiological dependence, which likely exceeded a mere exchange of amino acids, limited the growth of the entire consortium. Thus, the resulting metabolic and physiological intertwining decelerated growth and, as a consequence of this, also the ability of the whole consortium to adapt fast enough to a rapidly increasing environmental selection pressure. Another side effect of the harsh treatment was that these environmental conditions favoured mutants that reverted to metabolic autonomy, thus allowing them to escape the obligate relationship (Fig. 5). Under the conditions of our experiment, the cost of the interaction likely exceeded the benefits derived from it, thus selecting against obligate cooperation and favouring evolutionary independence. In fact, the evolutionary transition from an obligate mutualistic cooperation to a free-living state is one possibility of how mutualisms can break down. In phylogenetic analyses of obligate mutualistic interactions, free-living taxa are frequently nested within ancestrally mutualistic clades^38, 39, 40^. However, in many cases, the mechanistic explanation for the observed pattern remains elusive. Our study provides a possible answer: strong environmental selection pressures can challenge the ability of a obligately mutualistic consortium to adapt to changing environmental conditions, thus creating an incentive for mutants that are able to persist outside the interaction. Most likely, this mechanism is not restricted to microbial systems, but probably also operates in other types of mutualistic cooperation. An example is the interaction between reef corals and photosynthetic zooxanthellae, in which, due to the increasing pressure of global warming and ocean acidification, symbiotic algae are frequently lost from coral tissues (i.e. coral bleaching)^41, 42^. Moreover, the results of our work have also important ramifications for gut microbiota that are associated with humans or livestock. Studies have shown that these strongly interconnected gut microbiomes are essentially involved in a multitude of processes, including the harvesting of inaccessible nutrients as well as regulating the host’s intestinal homeostasis and immune responses^43, 44, 45^. Treating these microbial communities with antibiotics might therefore not only disturb their taxonomic composition, but also actively select against obligate (cooperative) metabolic interactions. Given the tremendous importance of these interactions for host health^43, 44, 45^, detrimental effects resulting from a loss of previously evolved metabolic interactions might impair host fitness and wellbeing for extended periods of time. Future work should assess how the treatment with antibiotics affects functional properties of gut microbiota and determine, whether potential side-effects can be neutralized by e.g. fecal transplants^46^.

Together, our results show that consortia engaging in an obligate mutualistic cooperation are significantly less able to respond to environmental selection pressures with evolutionary adaptation than physiologically autonomous individuals. Moreover, the resulting evolutionary cost of obligate cooperation can favour the emergence of genotypes that are capable of independent growth, thus destabilizing mutualistic interactions. These results significantly advance our understanding of the evolutionary consequences resulting from obligate cooperative interactions. Thus, these newly gained insights contribute to the development of a theoretical framework that can help to quantitatively predict the effects of anthropogenic alterations of natural ecosystems, ranging from climate change to the abuse of antibiotic drugs in clinical environments.

## Materials and Methods

### Strains and plasmids

*Escherichia coli* K12 BW25113 Δ*trpB ara*- Δ*LacZ* and *Escherichia coli* K12 BW25113 Δ*tyrA ara+ LacZ+* where used as experimental strains. Both auxotrophic strains were derived from a previous study, where they have been experimentally coevolved ^27^. This selection regime led to an enhanced production of the reciprocally exchanged amino acids at a cost to the producing cells. The kanamycin resistance cassette (i.e. neomycin phosphotransferase II (NPTII/Neo)) in the genome of both strains was removed using the plasmid pCP20 as described in ^47^.

### Culture conditions

All experiments were performed in 96 deep-well plates (maximal volume: 2 ml, Thermo Scientific Nunc) with minimal medium for *Azospirillium brasilense* (MMAB)^48^ without biotin and using glucose (5 g l^-1^) instead of sodium malate as a carbon source. All media components were purchased from VWR International GmbH (Darmstadt, Germany), Carl Roth GmbH & Co. KG (Karlsruhe, Germany), or Sigma-Aldrich (Darmstadt, Germany). Cultures were incubated at 30 °C under shaken conditions (200 rpm) for 72 hours. To enhance comparability and sustain growth, monocultures were supplemented with 100 mM of either tryptophan (Trp) or tyrosine (Tyr).

For plating, selective lysogeny broth (LB) agar plates and selective MMAB agar plates were used. These media were supplemented with 10 g l^-1^ of L-arabinose, 1 ml l^-1^ of 5% triphenyltetrazolium chloride (TTC), 1 mM of Isopropyl β-D-1-thiogalactopyranoside (IPTG), and 50 mg l^-1^ of 5-bromo-4-chloro-3-indolyl-β-D-galactopyranoside (x-Gal) to phenotypically distinguish differentially labelled strains in coculture.

### Evolution experiment

During the evolution experiment, monocultures of the tryptophan- and tyrosine-auxotrophic strains were simultaneously supplemented with both tryptophan and tyrosine (100 mM each). The media used for cocultivation did not contain any externally supplied amino acid. Each of the two monocultured strains involved in the evolution experiment started from one cryogenic stock. These strains were inoculated in 5 ml MMAB supplemented with both amino acids. After incubation for 72 h, liquid cultures were adjusted to an optical density at 600 nm (OD_600nm_) of 0.1 (SpectraMax microplate reader, Molecular Devices). Precultures were then used to inoculate 16 replicates for each of the three test cultures: (1) coculture of *E. coli* K12 BW25113 Δ*trpB ara*^*-*^ Δ*lacZ* and *E. coli* K12 BW25113 Δ*tyrA ara*^*+*^ *lacZ*^*+*^ (hereafter: CO), (2) monoculture of *E. coli* K12 BW25113 Δ*trpB ara*^*-*^ Δ*lacZ* (hereafter: TRP), and (3) monoculture of *E. coli* K12 BW25113 Δ*tyrA ara*^*+*^ *lacZ*^*+*^ (hereafter: TYR). In the case of both monocultures, 40 µl of preculture where used to inoculate 960 µl fresh MMAB containing both amino acids (100 mM each) and for cocultures, 20 µl of each auxotroph preculture of auxotrophs (TRP and TYR) was inoculated into 960 µl fresh MMAB without amino acids. 40 µl of the resulting cultures were transferred every 72 h into fresh medium. During the first two growth periods, no antibiotic treatment was applied to the cultures to allow populations to equilibrate and achieve homeostasis (see Supplementary Fig. 2). At the second transfer, each test culture was split up into five separate technical replicates, resulting in a total of 80 samples per treatment and strain combination. Simultaneously, the antibiotic treatment started at this step. Growth of all cultures was tracked by quantifying their population density (OD_600nm_), while propagating them to fresh medium. In addition, the death rate of all cultures was determined during the transfer: A culture was considered dead, when the OD_600nm_ value fell below the critical threshold of 0.01. In the course of the experiment, antibiotic concentrations were gradually increased (Fig. 1, Supplementary Fig. 3). Two main features characterized this ramping design. First, the antibiotic concentration doubled during each transfer until it reached the determined subminimal inhibitory concentration (sub-MIC, Supplementary Fig. 4), starting from the second transfer with one-eighth of the respective sub-MIC. Second, after reaching the sub-MIC level, antibiotic increment followed a linear function until it was slightly above the respective working concentrations. The rates, with which antibiotic concentrations were increased every transfer, were individually adjusted to each antibiotic, considering their sub-MIC values and targeted working concentrations. The corresponding working concentrations were 100 µg ml^-1^ for ampicillin, 25 µg ml^-1^ for chloramphenicol, 50 µg ml^-1^ for kanamycin, and 15 µg ml^-1^ for tetracycline. This information was provided by the manufacturer (Carl Roth GmbH & Co. KG, Karlsruhe, Germany) or derived from the Addgene archive^49^. These four antibiotics were chosen to maximise differences in terms of their effect (i.e. bacteriostatic or bactericidal), mode of action, and most frequently observed resistance mechanism^50, 51, 52^.

### Minimal inhibitory concentration (MIC)

To adjust the antibiotic pressure during the evolution experiment, sub-MIC values for cocultures were determined^53, 54^. Based on preliminary experiments (data not shown), the sub-MIC range was defined as 0.05 ± 0.005 OD_600nm_. Within this range, the bacterial culture is experiencing maximum antibiotic pressure, while maintaining sufficient growth to be transferred to the next growth cycle. Hereafter, this range is referred to as the *threshold zone*. An antibiotic concentration was considered as sub-MIC, if all measured OD_600nm_values at one antibiotic concentration fell within the threshold zone (Supplementary Fig. 4). In cases, in which OD_600nm_ values of more than one antibiotic concentration lay in the threshold zone, their average value was considered as sub-MIC. Moreover, if only a partial overlap of the values at one antibiotic concentration and the threshold zone was observed and the next measured distribution was not within the range, a mean value between those two was selected as sub-MIC. Only when there was no next measurement, the current concentration was picked. Based on these parameters, the sub-MIC values for the antibiotics used were 0.75 µg ml^-1^ for ampicillin, 3.25 µg ml^-1^ for chloramphenicol, 1.25 µg ml^-1^ for kanamycin, and 1.35 µg ml^-1^ for tetracycline (Supplementary Fig. 4). All MIC measurements were performed according to protocols of Wiegand *et al*. (2008) and Andrews (2001), using MMAB as growth medium. Growth depending on the current concentration of antibiotics was determined by quantifying population densities as OD_600nm_.

### Strain-level differences

An experiment was conducted to clarify how the two individual auxotrophic genotypes affected MIC values of the whole mutualistic consortium. To address this, the MIC of i) derived cooperative cocultures in both the presence and absence of the required amino acids, ii) the separated monocultures of the two genotypes within the coevolved cocultures, as well as iii) the auxotrophic genotypes that were cultivated as monocultures throughout the evolution experiment, was determined and compared among groups. The experiment was conducted with strains that evolved in the presence of either chloramphenicol or tetracycline (i.e. the two bacteriostatic antibiotics), because here a subset of replicated populations survived until the end of the main experiment (Fig. 2C,D).

### Separation of coevolved cocultures

Eight replicates from the chloramphenicol- and tetracycline-treated evolved cocultures were selected to separate individual auxotrophic genotypes from each other. For this, cultures were grown for 72 h and spread-plated on selective MMAB agar supplemented with tryptophan and tyrosine (100 mM each). Separation was performed by picking single colonies from these plates and streaking them on five different selective plates (MMAB+Trp+Tyr, MMAB, MMAB+Trp, MMAB+Tyr, LB), to determine their phenotype. From chloramphenicol-treated cocultures, three colonies belonging to each genotype (TRP and TYR) were isolated per replicate. However, for tetracycline-treated cocultures, only three colonies could be isolated that showed the tryptophan background. This was most likely due to the low abundance of tryptophan-auxotrophic genotypes in these cultures (Fig. 5).

### Effect of coevolutionary history and metabolic dependence

An experiment was conducted to determine the interactive effect between (i) a previous coevolutionary history in a given stressful environment as well as (ii) the metabolic dependence on another genotype on the ability of strains to cope with environmental stress. For this, strains that evolved in the presence of chloramphenicol and tetracycline either alone (i.e., monoevolved) or in coculture were grown as coculture (both consisting of either monoevolved or coevolved strains, Supplementary Table 4) in the presence and absence of the two required amino acids (100 mM each). In total, eight randomly selected pairs of mono- or coevolved tryptophan and tyrosine auxotrophs were used, which have been isolated from the 15^th^ transfer of the main experiment. All of these populations were cultivated at different concentrations of the antibiotics chloramphenicol and tetracycline according to previous experiments (data not shown) and the population densities (OD_600nm_) they achieved at 72 h post-inoculation were quantified.

### Phenotypic reversion

To quantify the rate of phenotypic reversion within cultures that evolved in presence of chloramphenicol or tetracycline, 15 replicates from the 15^th^ transfer of each test culture type (CO, TRP, and TYR) were randomly selected to determine whether the population was still auxotrophic or if also prototrophic revertants have arisen to detectable levels. Precultures of evolved monocultures were supplemented with both tryptophan and tyrosine (100 mM each), while precultures of coevolved cocultures were grown in the absence of amino acids for 72 h in liquid MMAB. In addition, all cultures were supplemented with either chloramphenicol (13.75 µg ml^-1^) or tetracycline (8.25 µg ml^-1^) to mimic the conditions of the evolution experiment as closely as possible, while maintaining sufficient growth. These precultures were spread-plated on selective LB agar and single colonies were picked from each test sample and streaked on four different selective plates (MMAB, MMAB + Trp, MMAB + Tyr, LB). Colonies that grew on all plates comparably well were considered as phenotypic revertants, based on the apparent loss of their auxotrophic phenotype. In this way, it was possible to differentiate and classify auxotrophic and prototrophic colonies within cocultures with a lower detection limit of 2.5 x 10^4^ CFU ml ^-1^ and an upper detection limit of 2.5 x 10^6^ CFU ml ^-1^.

### Statistical analysis

An unsupervised learning algorithm was used to characterize the evolutionary trajectory followed by mono- and cocultures of auxotrophs during the evolution experiment. In particular, profiles of cell densities between cultures and across transfers were compared using a variational Gaussian mixture algorithm to detect clusters in the data. To compare survival distributions between cultures of this experiment, log-rank tests were calculated for each pair of cultures.

Differences in the evolvability of coevolved cocultures and individually evolved strains were assessed using linear mixed models with bacterial growth as the response variable. To compare strain types in the presence and absence of amino acids, contrasts between their growth levels were calculated using the *emmeans* package function *emmeans* from R^55^ (Supplementary Information 1).

All remaining statistical tests were performed in SPSS (Version 25, IBM^®^). Differences in the level of antibiotic resistance between coevolved and monoevolved strains were identified using an ANOVA followed by a Tamhane’s post-hoc test. Significant differences in the population-level ratio in the number of phenotypic reversion of evolved monocultures and cocultures were identified using a chi-square test. Normal distribution of datasets was assessed using the Kolmogorov-Smirnov test and homogeneity of variances was determined by applying Levene’s test. Variances were considered to be homogeneous when P > 0.05.

## Acknowledgements

The authors thank Daniel Preußger for providing strains and advice on experiments. This manuscript benefitted greatly from discussions with Piyali Pal Chowdhury, Samir Giri, and all other members of the Kostlab. This work was funded by the German Research Foundation (SFB 944, P19: CK; SPP1617, KO 3909/2-1: CK), the International Graduate School *EvoCell* (CK), and the University of Osnabrück (LO).

## Author contributions

Conceived the study: C.K., B.P.

Designed the experiments: B.P., C.K., L.O.

Performed the experiments: B.P., M.H.

Analyzed the results: L.O., B.P.

Interpreted the results: B.P., L.O., C.K.

Wrote the manuscript: B.P., C.K

Amended the manuscript: L.O.

