## Supplemental Material for "Obligate mutualistic cooperation limits evolvability"

**Table of contents** **Page**

Supplementary Figures……………………………………………………...………................... 2

Supplementary Tables…….…………………………………………………….………………..…. 6

Supplementary Information.…...……………………………………..…………………..………… 11

Supplementary References……………………………………….…………………..……………. 13

### Supplementary Figures

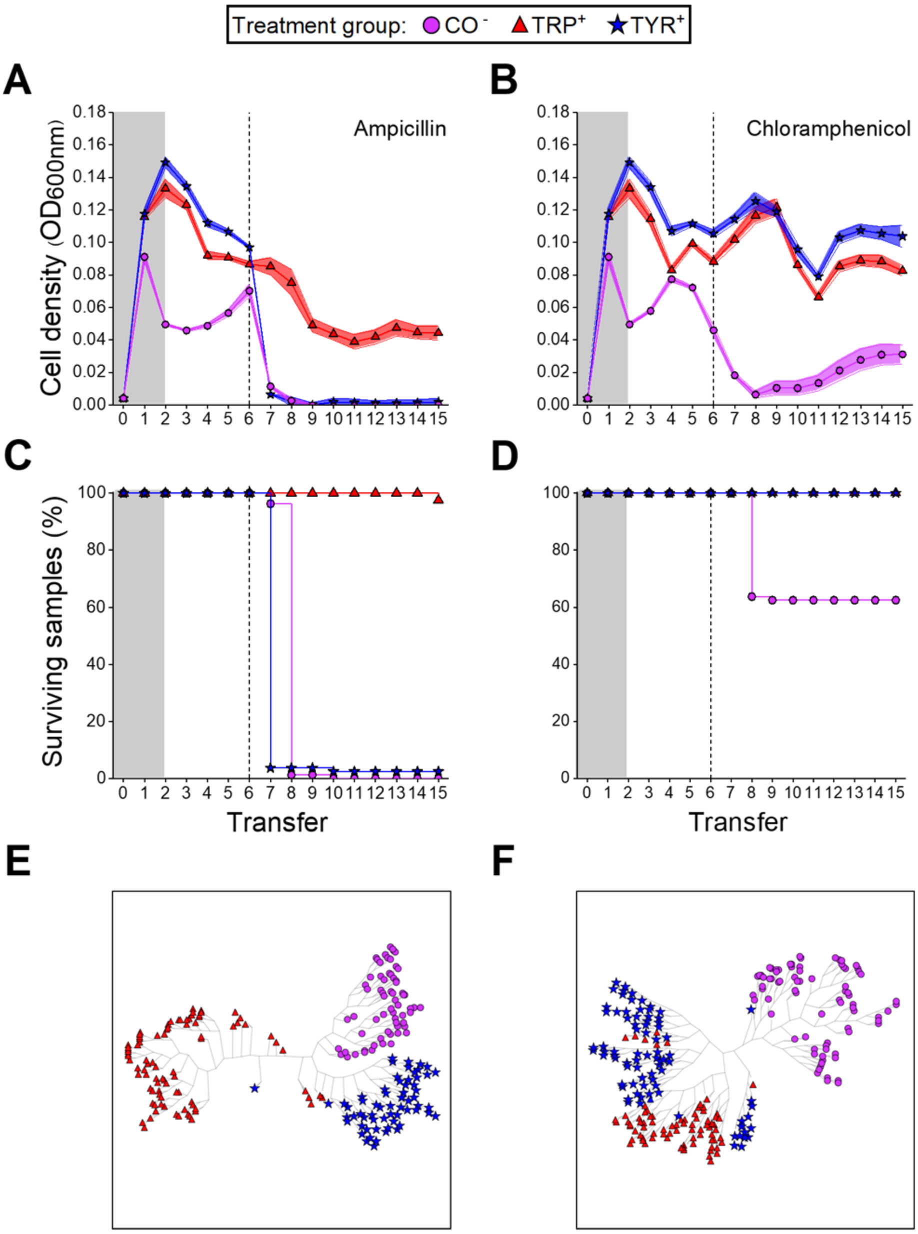


**Supplemental Figure 1 | Data of the ampicillin and chloramphenicol treatment of the evolution experiment (Fig. 2). (A, B)** Mean growth (±95% confidence interval, n = 80 per point) quantified as $OD_{600nm}$ and **(C, D)** proportion of surviving replicates in percent (n = 80 per strain) of auxotrophic monocultures (TRP, TYR) and mutualistic cocultures (CO) of the tryptophan (TRP) and tyrosine (TYR) auxotrophic strains over the course of the evolution experiment. Antibiotic concentrations were increased in a stepwise manner after each transfer (i.e. every 72 h) (Supplementary Fig. 3). Grey-shaded areas indicate periods without antibiotic treatment. **(A, C, E)** ampicillin treatment **(B, D, F)** chloramphenicol treatment. Dashed lines mark the point, at which antibiotic concentrations exceed sub-MIC values. Monocultures were supplemented with both amino acids (100 mM each), while cocultured bacteria depended on the amino acids provided by their respective cross-feeding partner. **(E, F)** Clustering trees of cell density profiles of the different cultures across transfers in the evolution experiment. Each leaf within a given tree represents a replicate (n = 80 per strain). A radial embedding layout was used to display trees.

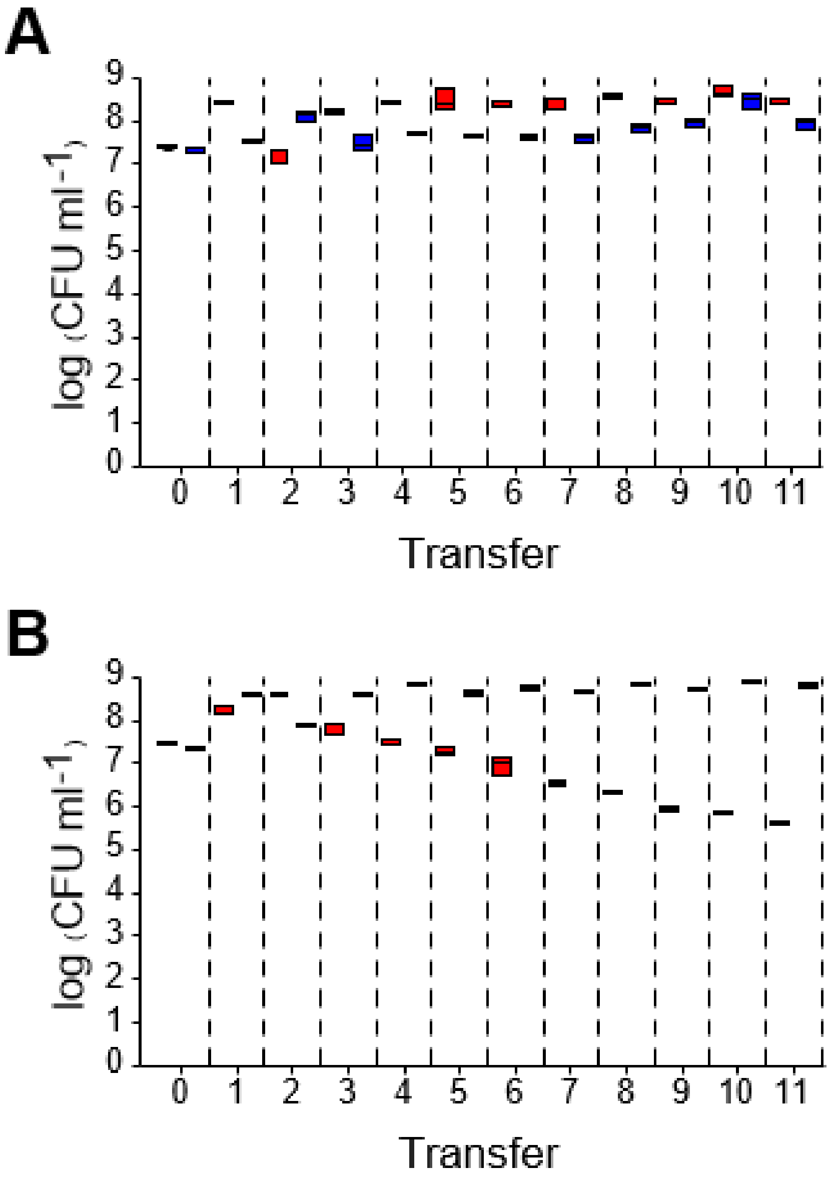


**Supplemental Figure 2 | Cocultures of auxotrophs reach a stable equilibrium in the absence, but not the presence of externally provided amino acids.** Cocultures of *E. coli* BW25113 *ΔtrpB* ara$^-$ Δ*lacZ* (TRP, red boxes) and *E. coli* BW25113 *ΔtyrA* ara$^+$ lacZ$^+$ (TYR, blue boxes) were serially propagated every 72 h **(A)** without and **(B)** with amino acid supplementation (100 mM each). Shown are the numbers of colony-forming units (CFU) of each strain per millilitre of culture. Boxes show median values (horizontal line in boxes) and the upper and lower quartiles (i.e. 25-75% of data, boxes). Whiskers indicate the 1.5x interquartile range (n = 4).

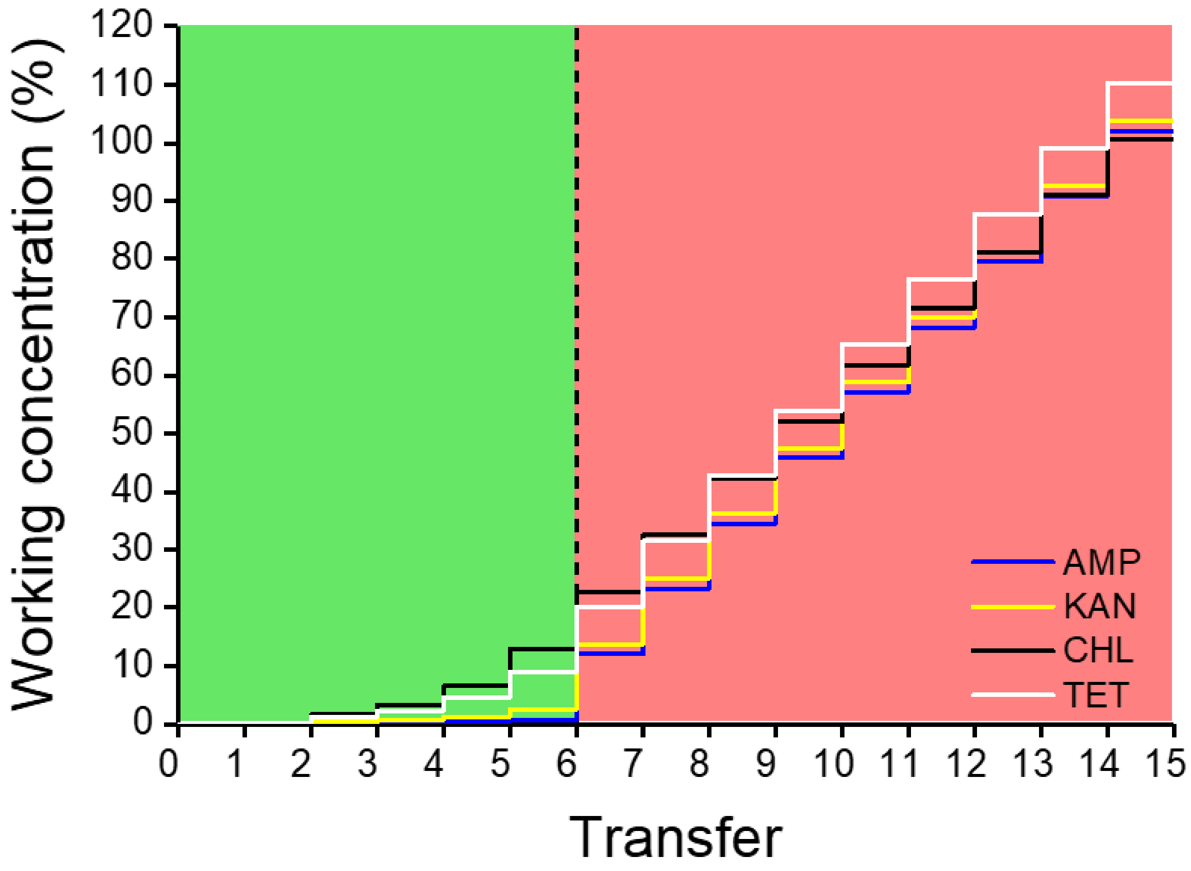


**Supplemental Figure 3 | Ramping of antibiotic concentrations during the evolution experiment.** Concentrations of antibiotics displayed in percent of the respective working concentration (i.e. ampicillin: 100 µg $ml^{-1}$, kanamycin: 50 µg $ml^{-1}$, chloramphenicol: 25 µg $ml^{-1}$, and tetracycline: 15 µg $ml^{-1}$). The green area represents antibiotic concentrations below and the red area above the determined sub-minimal inhibitory concentrations (sub-MIC). Antibiotic concentrations doubled at each transfer, starting from the second transfer. Cultures were transferred every 72 h to fresh medium.

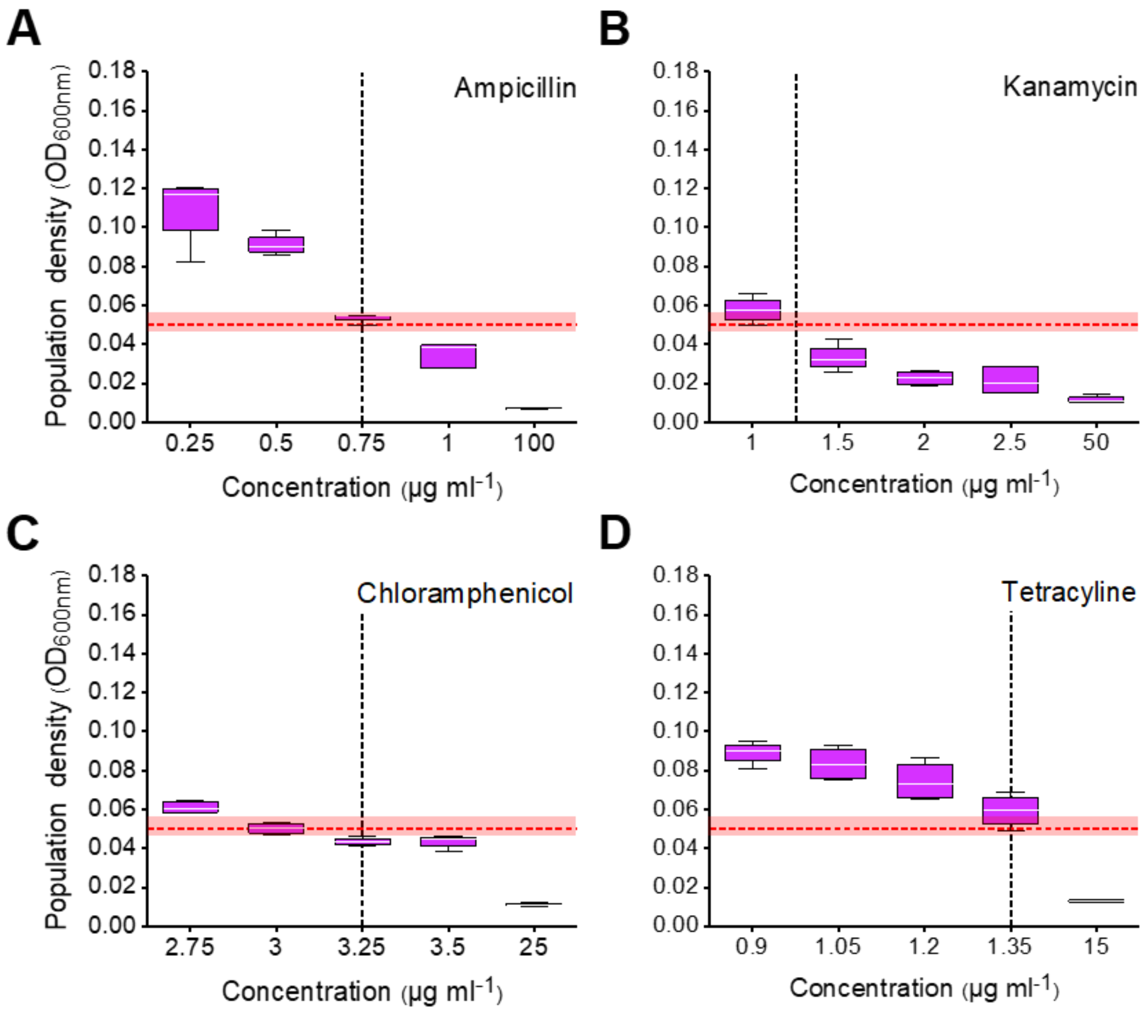


**Supplemental Figure 4 | Determination of sub-minimal inhibitory concentrations (sub-MIC) for populations of cocultured auxotrophs.** Cocultures of *E. coli* BW25113 *ΔtrpB* ara$^-$ Δ*lacZ* and *E. coli* BW 25113 *ΔtyrA* ara$^+$ lacZ$^+$ (purple boxes) were grown in minimal medium supplemented with increasing concentrations of the antibiotic **(A)** ampicillin, **(B)** kanamycin, **(C)** chloramphenicol, and **(D)** tetracycline. Growth was evaluated by measuring the optical density ($OD_{600nm}$) of cultures. The red dashed line and the red-shaded area represent the threshold zone (0.05 ± 0.005 $OD_{600nm}$). The black dashed line indicates the sub-MIC. Boxes show median values (horizontal line in boxes) and the upper and lower quartiles (i.e. 25-75% of data, boxes). Whiskers indicate the 1.5x interquartile range (n = 4).

### Supplemental Tables

**Supplemental Table 1 | Log-rank test results of the survival analysis.** Dashes indicate comparisons with no measurable difference due to the fact that all replicates within both groups survived until the end of the evolution experiment. See Supplemental Table 4 for abbreviations of experimental groups.

| Figure | Treatment | Group 1 | Group 2 | P | $\chi^2$ | n |
|---|---|---|---|---|---|---|
| 2 C | Kanamycin | CO | TRP | $1.3 * 10^{-33}$ | 145.946 | 80 |
| 2 C | Kanamycin | CO | TYR | $2.1 * 10^{-27}$ | 117.6 | 80 |
| 2 C | Kanamycin | TRP | TYR | 0.16 | 1.97 | 80 |
| 2 D | Tetracycline | CO | TRP | $1.9 * 10^{-16}$ | 67.71 | 80 |
| 2 D | Tetracycline | CO | TYR | $1.9 * 10^{-16}$ | 67.71 | 80 |
| 2 D | Tetracycline | TRP | TYR | - | - | 80 |
| Sup. 4 C | Ampicillin | CO | TRP | $1.1 * 10^{-36}$ | 160.023 | 80 |
| Sup. 4 C | Ampicillin | CO | TYR | $2.3 * 10^{-25}$ | 108.338 | 80 |
| Sup. 4 C | Ampicillin | TRP | TYR | $3.2 * 10^{-35}$ | 153.352 | 80 |
| Sup. 4 D | Chloramphenicol | CO | TRP | $1.3 * 10^{-9}$ | 36.76 | 80 |
| Sup. 4 D | Chloramphenicol | CO | TYR | $1.3 * 10^{-9}$ | 36.76 | 80 |
| Sup. 4 D | Chloramphenicol | TRP | TYR | - | - | 80 |

**Supplemental Table 2 | Statistical results of comparisons shown in figure 3.** Experimental cultures were classified as coevolved ($_{CO}$), monoevolved ($_{MO}$), amino acid supplemented ($^{+}$) or unsupplemented ($^{-}$). For abbreviations of experimental groups see Supplemental Table 4.

| Figure | Group 1 | Group 2 | P | F | n |
|---|---|---|---|---|---|
| 3 A | $CO^-$ | $CO^+$ | 1.000 | 33.546 | 8 |
| 3 A | $CO^-$ | $TRP_{CO}^+$ | 0.006 | 33.546 | 8 |
| 3 A | $CO^-$ | $TYR_{CO}^+$ | 0.005 | 33.546 | 8 |
| 3 A | $CO^-$ | $TRP_{MO}^+$ | 0.016 | 33.546 | 8 |
| 3 A | $CO^-$ | $TYR_{MO}^+$ | < 0.001 | 33.546 | 8 |
| 3 A | $CO^+$ | $TRP_{CO}^+$ | 0.557 | 33.546 | 8 |
| 3 A | $CO^+$ | $TYR_{CO}^+$ | 0.502 | 33.546 | 8 |
| 3 A | $CO^+$ | $TRP_{MO}^+$ | 0.035 | 33.546 | 8 |
| 3 A | $CO^+$ | $TYR_{MO}^+$ | 0.002 | 33.546 | 8 |
| 3 A | $TRP_{CO}^+$ | $TYR_{CO}^+$ | 1.000 | 33.546 | 8 |
| 3 A | $TRP_{CO}^+$ | $TRP_{MO}^+$ | < 0.001 | 33.546 | 8 |
| 3 A | $TRP_{CO}^+$ | $TYR_{MO}^+$ | < 0.001 | 33.546 | 8 |
| 3 A | $TYR_{CO}^+$ | $TRP_{MO}^+$ | < 0.001 | 33.546 | 8 |
| 3 A | $TYR_{CO}^+$ | $TYR_{MO}^+$ | < 0.001 | 33.546 | 8 |
| 3 A | $TRP_{MO}^+$ | $TYR_{MO}^+$ | 0.936 | 33.546 | 8 |
| 3 B | $CO^-$ | $CO^+$ | 0.993 | 240.555 | 8 |
| 3 B | $CO^-$ | $TRP_{CO}^+$ | < 0.001 | 240.555 | 8 |
| 3 B | $CO^-$ | $TYR_{CO}^+$ | 0.016 | 240.555 | 8 |
| 3 B | $CO^-$ | $TRP_{MO}^+$ | < 0.001 | 240.555 | 8 |
| 3 B | $CO^-$ | $TYR_{MO}^+$ | 0.993 | 240.555 | 8 |
| 3 B | $CO^+$ | $TRP_{CO}^+$ | < 0.001 | 240.555 | 8 |

| | | | | | |
|---|---|---|---|---|---|
| 3 B | $CO^{+}$ | $TYR_{CO}^{+}$ | 0.010 | 240.555 | 8 |
| 3 B | $CO^{+}$ | $TRP_{MO}^{+}$ | < 0.001 | 240.555 | 8 |
| 3 B | $CO^{+}$ | $TYR_{MO}^{+}$ | 1.000 | 240.555 | 8 |
| 3 B | $TRP_{CO}^{+}$ | $TYR_{CO}^{+}$ | < 0.001 | 240.555 | 8 |
| 3 B | $TRP_{CO}^{+}$ | $TRP_{MO}^{+}$ | 0.049 | 240.555 | 8 |
| 3 B | $TRP_{CO}^{+}$ | $TYR_{MO}^{+}$ | < 0.001 | 240.555 | 8 |
| 3 B | $TYR_{CO}^{+}$ | $TRP_{MO}^{+}$ | < 0.001 | 240.555 | 8 |
| 3 B | $TYR_{CO}^{+}$ | $TYR_{MO}^{+}$ | 0.010 | 240.555 | 8 |
| 3 B | $TRP_{MO}^{+}$ | $TYR_{MO}^{+}$ | < 0.001 | 240.555 | 8 |

**Supplemental Table 3 | Statistical results of comparisons shown in figure 4.** Experimental cultures were classified as amino acid supplemented ($^+$) or unsupplemented ($^-$). For abbreviations of experimental groups see Supplemental Table 4.

| Figure | Treatment | Strain 1 | Strain 2 | P | t ratio | df |
|---|---|---|---|---|---|---|
| 4 A | Chloramphenicol | $CO_{EVO}^-$ | $CO_{AUX}^-$ | < 0.001 | - 5.715 | 21.4 |
| 4 B | Chloramphenicol | $CO_{EVO}^+$ | $CO_{AUX}^+$ | < 0.001 | - 7.418 | 21.4 |
| 4 A,B | Chloramphenicol | $CO_{EVO}^-$ | $CO_{EVO}^+$ | 0.0675 | 2.637 | 21.4 |
| 4 A,B | Chloramphenicol | $CO_{AUX}^-$ | $CO_{AUX}^+$ | < 0.001 | - 15.928 | 21.4 |
| 4 C | Tetracycline | $CO_{EVO}^-$ | $CO_{AUX}^-$ | < 0.001 | - 4.311 | 28.9 |
| 4 D | Tetracycline | $CO_{EVO}^+$ | $CO_{AUX}^+$ | < 0.001 | - 7.661 | 36.9 |
| 4 C,D | Tetracycline | $CO_{EVO}^-$ | $CO_{EVO}^+$ | 0.7452 | - 1.009 | 28.9 |
| 4 C,D | Tetracycline | $CO_{AUX}^-$ | $CO_{AUX}^+$ | < 0.001 | - 11.306 | 36.9 |

**Supplemental Table 4 | Strains and abbreviations.**

| **Abbreviation** | **Strain** |
|---|---|
| TRP | *E. coli* BW25113 Δ*trpB* ara- Δ*lacZ* |
| TYR | *E. coli* BW 25113 Δ*tyrA* ara$^{+}$ lacZ$^{+}$ |
| CO | *E. coli* BW25113 Δ*trpB* ara$^{-}$ Δ*lacZ* cocultured with *E. coli* BW 25113 Δ*tyrA* ara$^{+}$ *lacZ*$^{+}$ |
| $CO_{EVO}$ | coevolved *E. coli* BW25113 Δ*trpB* ara$^{-}$ Δ*lacZ* and *E. coli* BW 25113 Δ*tyrA* ara$^{+}$ *lacZ*$^{+}$ |
| $CO_{AUX}$ | monoevolved *E. coli* BW25113 Δ*trpB* ara$^{-}$ Δ*lacZ* and *E. coli* BW 25113 Δ*tyrA* ara$^{+}$ lacZ$^{+}$ in coculture |

**Supplemental Information 1**

**Statistical model used to analyse growth patterns during the evolution experiment**

Linear mixed models (LMM) were used with *bacterial growth* as the response variable. As the antibiotics chloramphenicol and tetracycline were applied in different concentrations (depending on their respective MIC, Fig. 4), two separate analyses were conducted: one for chloramphenicol and one for tetracycline (for each case: n = 320). As fixed effects, the *antibiotic concentration* and the *strain type in the presence or absence of amino acids* were included. We controlled for multiple measurements by including replicates as a random effect in the model.

Before running the models, the two numerical covariates were z-transformed to a mean of zero and a standard deviation of one, and the response variable were square root-transformed to achieve a more symmetrical distribution. Models were fitted in R[1] by using the function *lmer* from the R package lme4[2]. The LMMs were verified to test whether the assumptions of normal distribution and homogeneous residuals were fulfilled by visually inspecting a qqplot and plotting the residuals against the fitted values. In both models, no obvious deviation from the abovementioned assumptions were detected. Additionally, model stability was scrutinized by excluding each level of the random effect at a time from the data. A comparison of the model estimates derived for the reduced data set with those derived by the full data set ruled out the existence of overly influential cases. Variance Inflation Factors were calculated (VIF,[3]) using the function *vif* of the R-package car[4]. The results did not indicate collinearity to be an issue (largest VIF = 2.00). For LMMs, statistical significance of the full model was determined by comparing its fit with that of the null model comprising only the random effect, using a likelihood ratio test (LRT)[5]. For this, the R function *anova package stats* was used with argument test set to *Chisq*. The P-values for fixed effects were based on a likelihood ratio test, comparing the full model with a reduced model excluding the fixed effects[6, 7] using the R function *drop1* with argument test set to *Chisq*. To allow for an LRT, the models were fitted using Maximum Likelihood (rather than Restricted Maximum Likelihood;[8]).

The results of the tests evaluating the models proposed revealed a clear influence of the fixed effects on the response variable (i.e. bacterial growth). LRT comparing full and null model for chloramphenicol: $\chi^2 = 93.999$, df = 7, P < 0.001 and tetracycline: $\chi^2 = 88.155$, df = 7, P < 0.001. Each fixed factor: a) antibiotic concentration and b) strain type in the presence or absence of amino acids, has a statistically significant effect on bacterial growth. For chloramphenicol, a: LRT = 24.185, P < 0.001; b: LRT = 77.625, P < 0.001. For tetracycline, a: LRT = 24.838, P < 0.001; b: LRT = 48.882, P < 0.001.

To compare between strain types in the presence or absence of amino acids, contrasts between growth levels were calculated using *emmeans* from the emmeans package function in R[9].
